# The SARS-CoV-2 cytopathic effect is blocked with autophagy modulators

**DOI:** 10.1101/2020.05.16.091520

**Authors:** Kirill Gorshkov, Catherine Z. Chen, Robert Bostwick, Lynn Rasmussen, Miao Xu, Manisha Pradhan, Bruce Nguyen Tran, Wei Zhu, Khalida Shamim, Wenwei Huang, Xin Hu, Min Shen, Carleen Klumpp-Thomas, Zina Itkin, Paul Shinn, Anton Simeonov, Sam Michael, Matthew D. Hall, Donald C. Lo, Wei Zheng

**Author notes:** Correspondence: Kirill Gorshkov –, Wei Zheng –.

## Abstract

SARS-CoV-2 is a new type of coronavirus capable of rapid transmission and causing severe clinical symptoms; much of which has unknown biological etiology. It has prompted researchers to rapidly mobilize their efforts towards identifying and developing anti-viral therapeutics and vaccines. Discovering and understanding the virus’ pathways of infection, host-protein interactions, and cytopathic effects will greatly aid in the design of new therapeutics to treat COVID-19. While it is known that chloroquine and hydroxychloroquine, extensively explored as clinical agents for COVID-19, have multiple cellular effects including inhibiting autophagy, there are also dose-limiting toxicities in patients that make clearly establishing their potential mechanisms-of-action problematic. Therefore, we evaluated a range of other autophagy modulators to identify an alternative autophagy-based drug repurposing opportunity. In this work, we found that 6 of these compounds blocked the cytopathic effect of SARS-CoV-2 in Vero-E6 cells with EC_50_ values ranging from 2.0 to 13 μM and selectivity indices ranging from 1.5 to >10-fold. Immunofluorescence staining for LC3B and LysoTracker dye staining assays in several cell lines indicated their potency and efficacy for inhibiting autophagy correlated with the measurements in the SARS-CoV-2 cytopathic effect assay. Our data suggest that autophagy pathways could be targeted to combat SARS-CoV-2 infections and become an important component of drug combination therapies to improve the treatment outcomes for COVID-19.

## Introduction

The COVID-19 global viral pandemic caused by SARS-CoV-2 began in late 2019 originating from Wuhan, Hubei Province, China (*1*). SARS-CoV-2 is a member of the *Coronaviridae* family of positive single-stranded RNA viruses. As of May 14, 2020, there have been over 4,405,000 infections worldwide and 300,000 deaths (*2*). While not the deadliest virus in the past century, it is highly infectious (estimated R_0_ = 5.7) (*3*). The absolute number of infections and mortality will not be known for several years (*4*).

SARS-CoV-2 infection in humans produces a disease called <u>co</u>rona<u>v</u>irus <u>d</u>isease of 2019, COVID-19 (*5*) (*6*). It is related to the 2003 coronavirus outbreak of SARS-CoV, the original SARS. Fortunately, the virus was well-contained and no cases have been reported since 2004. COVID-19 symptoms range from mild fever, tiredness, and dry cough, to acute respiratory distress syndrome, stroke due to blood clots, cardiac and renal damage, and death (*7*). While some clinical symptoms are common among patients with severe disease, its epidemiology and the mechanisms of disease pathology are still unclear and need to be further studied.

The research and clinical responses have been unprecedented, and much of the effort is focused on identifying therapeutics, including drug repurposing efforts with the experimental anti-Ebola virus drug remdesivir (*8, 9*) and developing vaccines. Chloroquine (CQ), an older FDA-approved anti-malarial drug, along with its better tolerated analog hydroxychloroquine (HCQ), have been reported to inhibit SARS-CoV-2 infection *in vitro* and show some promise in patients (*10–12*). In mice, CQ and HCQ display antiviral effects against human coronavirus strain OC43 (*13*) human enterovirus EV71 (*14*), Zika virus (*15*), and human influenza virus H5N1 (*16*) CQ was not effective in reducing viral titers in the lungs of mice infected with SARS-CoV, although it did induce a reduction in markers of inflammation (*17*). The efficacy of CQ in animal models of SARS-CoV-2 has not yet been reported. CQ and HCQ have been reported to elicit antiviral activity via a number of mechanisms of action including modulation of autophagy.

Autophagy maintains cellular organelle homeostasis by clearing cellular waste and providing the cell with a supply of energy when nutrients are scarce (*18*). Autophagy also functions as the first line of defense to cleanse the cell of invading pathogens such as viruses, and plays an important role in mediating the innate immune response (*19*). The activation of autophagy engulfs virions inside host cells via the formation of autophagosomes that subsequently fuse with acidic lysosomes to form autolysosomes through a pH-dependent mechanism. The autolysosomal contents are then degraded by the lysosomal hydrolases. This entire autophagy cycle is called autophagic flux and plays a key role in processing invading viruses. In Drosophila, for example, NF-kB activation during Zika virus infection leads to elevated levels of Drosophila stimulator of interferon genes and increased autophagy in the brain (*20*). Unfortunately, some viruses have developed mechanisms to escape autophagy (*21*), avoid the immune response (*22*), and hijack the autophagosomes for viral replication (*23, 24*).

Viral hijacking of the endocytic pathway for viral entry and utilization of the autophagic machinery for their replication has been reported (*24, 25*). However, some conflicting data has demonstrated that SARS-CoV replication is independent of autophagic mechanisms (*26*). Viruses have also evolved strategies for escaping degradation through the inhibition of autophagosome-lysosome fusion and autophagic flux (*27, 28*). Nonetheless, given that autophagy inhibitors may act on multiple points within the viral life cycle, treating infection with lysosomotropic compounds may be a viable strategy for suppressing viral attack, and explain the potential therapeutic benefits of CQ and HCQ that have been reported in COVID-19 (*29*).

Autophagy inhibitors including CQ, HCQ, and others with related chemical structures are known to prevent the fusion of autophagosomes and lysosomes and blocks the later stages of autophagic flux (*30*). CQ, in addition to its inhibitory effects on autophagy, has been reported to have broad antiviral effects through several mechanisms of action. One in particular is the disruption of the early steps in the viral life cycle including the release of the virus from the endosome when endocytosis is used for viral entry (*31, 32*). Given the fact that the basic properties of CQ and similar molecules lead to their accumulation in acidic compartments and raise their pH, viruses that depend on low acidic pH for entry, uncoating, and replication can no longer execute these functions. While they exert multiple cellular effects, their characterized inhibition of autophagic flux and elevation of vesicular pH is consistent with the antiviral efficacy *in vitro* (*33*).

Accordingly, a recent SARS-CoV-2 study by Liu *et al*. has proposed that these drugs may act by preventing the progression of the virions through the endocytic pathway, thereby inhibiting release of the viral genome (*12*).

In this study, we have identified 6 known autophagy inhibitors that reduce the cytopathic effect (CPE) of SARS-CoV-2 in Vero-E6 cells. We have further investigated the effects of the compounds on markers of autophagy in several different cell lines using LC3B autophagic marker immunostaining as well as LysoTracker dye staining (*34*). These assays revealed comparable potency of the compounds for inhibiting autophagy compared to inhibition of the virally-induced cytopathic effect. Altogether, we demonstrate that the autophagy inhibitors effectively inhibited SARS-CoV-2 infection *in vitro*, indicating autophagy as a viable target pathway for COVID-19 drug discovery.

## Results

### Autophagy inhibitors block the cytopathic effect of SARS-CoV-2

We employed a cell-based assay using Vero-E6 host cells that measures the CPE of SARS-CoV-2 (Fig. 1).

**Fig. 1.**
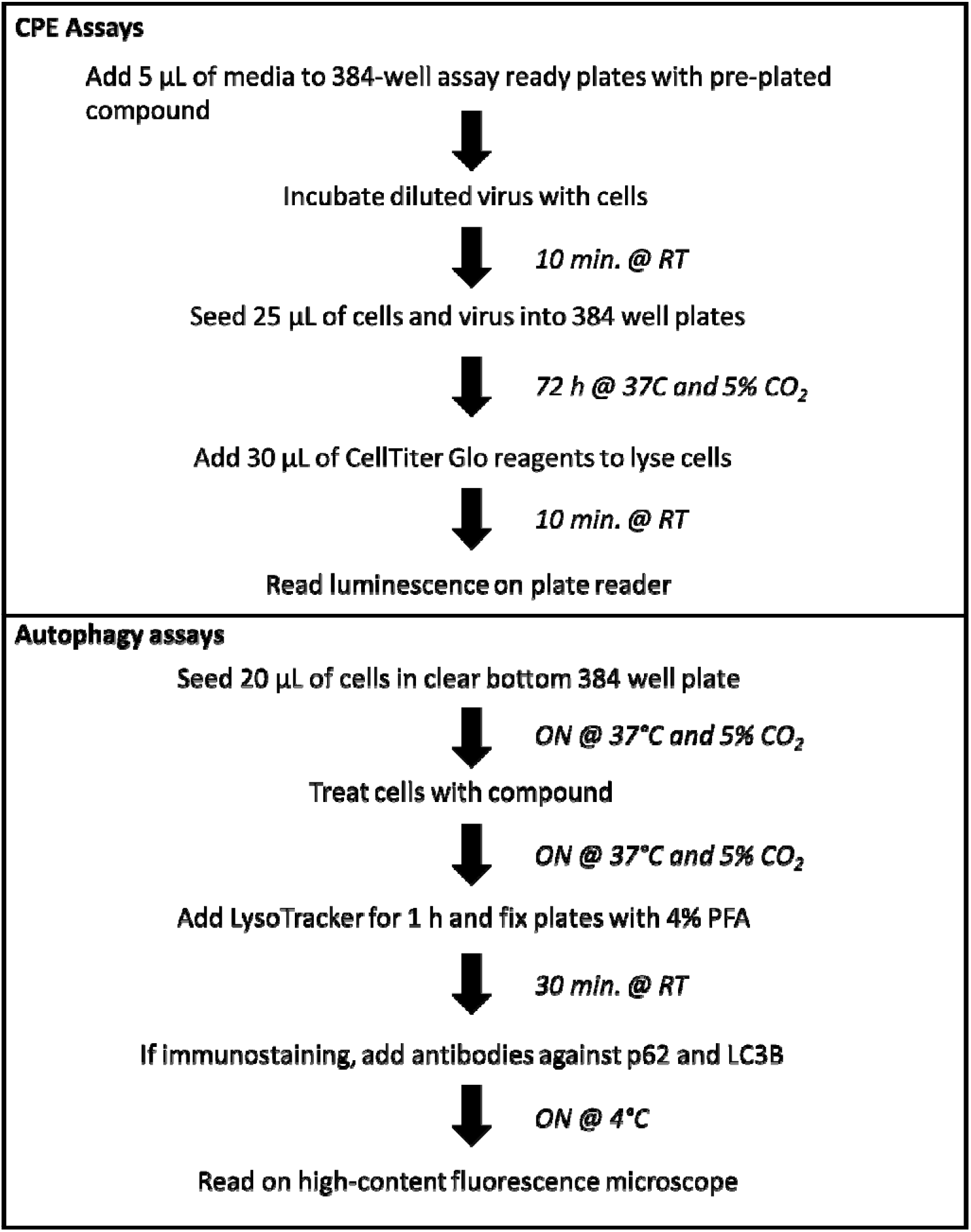
Workflow overview for CPE assay. Activities and incubation times are shown in a workflow.

The CPE reduction assay is a widely-employed assay format to screen for antiviral agents, and it can be scaled for high-throughput screening (*35, 36*). In this assay, host cell death is a consequence of the viral infection and cell viability is a surrogate readout for viral infection that can be measured with a range of cell viability assays. All compounds were tested in doseresponse, and ‘hit’ antiviral compounds are those that protect the host cells from viral CPE. To increase infectivity of SARS-CoV-2 in the CPE assay, we used a clone of Vero-E6 that had previously been selected for high ACE2 expression (*35*). The cell viability measurements were normalized to cells not infected with the virus (100% activity) and untreated cells infected with virus (0% activity; virus completely kills cells). As a counter-assay, all compounds were tested against cells not exposed to virus, in order to identify compounds that exerted cytotoxicity against Vero E6 cells.

Given that autophagy inhibitors including HCQ have shown efficacy against many different types of viruses (*31*) including SARS-CoV-2 in CPE assays (*12*), we assessed the protective effect of a group of autophagy inhibitors including ROC-325, clomipramine, hycanthone, verteporfin, CQ, HCQ, and mefloquine in 384-well plates (Fig. 2).

**Fig. 2.**
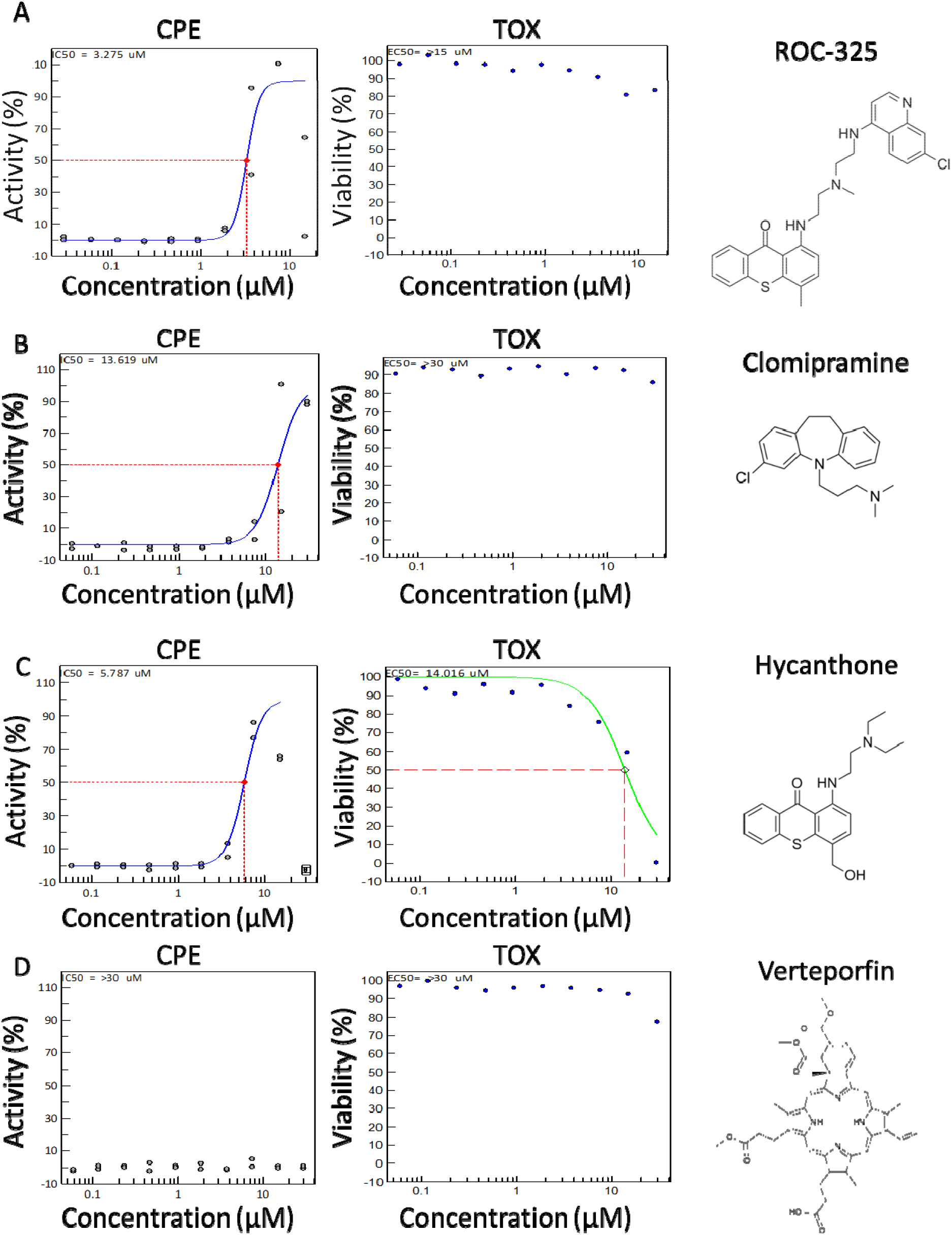
CPE activity and Toxicity for ROC-325, clomipramine, hycanthone, and verteporfin. **(A)** ROC-325**, (B)** clomipramine, **(C)** hycanthone, and **(D)** verteporfin CPE activity (blue curve, left graph) and Toxicity (green curve, right graph) 10 point, 1:2 dilution concentration-response curves starting at 30.0 μM down to 2.29 nM, along with their structure. ROC-325 started at 15 μM down to 1.14 nM. Red dashed line indicates EC50 or CC50 for CPE and Toxicity assays, respectively. Duplicate values shown for each concentration. Curves generated using non-linear regression.

While CQ was the most potent compound (discussed below), ROC-325 was the second most potent with an EC50 of 3.28 μM and less than 20% cytotoxicity at 30.0 μM (Fig. 2A), indicating a greater than 10-fold selectivity index (SI) between antiviral and cytotoxic concentrations. Clomipramine exhibited an EC50 of 13.6 μM while inducing less than 20% cytotoxicity at 30.0 μM (Fig. 2B). Hycanthone demonstrated an EC50 of 5.79 μM and a cytotoxicity CC50 of 14.0 μM (Fig. 2C). Hycanthone’s concentration-response was bell-shaped due to reduction of cell viability by almost 100% at 30 μM. Verteporfin was inactive in the screen against SARS-CoV-2 CPE and reduced cell viability by approximately 22% at 30.0 μM (Fig. 2D).

The anti-malarial drugs CQ and HCQ inhibited viral CPE with an EC50 of 2.01 μM and 4.47 μM, respectively, with no associated cell toxicity (Fig. 3A,B). HCQ was the third most potent compound tested in the CPE. Mefloquine, a related anti-malarial compound, exhibited an EC50 of 3.85 μM with an associated cell toxicity CC50 of 8.78 μM and 100% cytotoxicity at 15.0 to 30.0 μM (Fig. 3C). For comparison, remdesivir, the nucleotide analog inhibitor of RNA-dependent RNA polymerase for a number of viruses and top clinical candidate for SARS-CoV-2 (*8, 9, 37*), exhibited an EC50 of 7.04 μM with no apparent cytotoxicity (Fig. 3D). The EC50 values for all of the autophagy inhibitor compounds are summarized in Table 1.

**Fig. 3.**
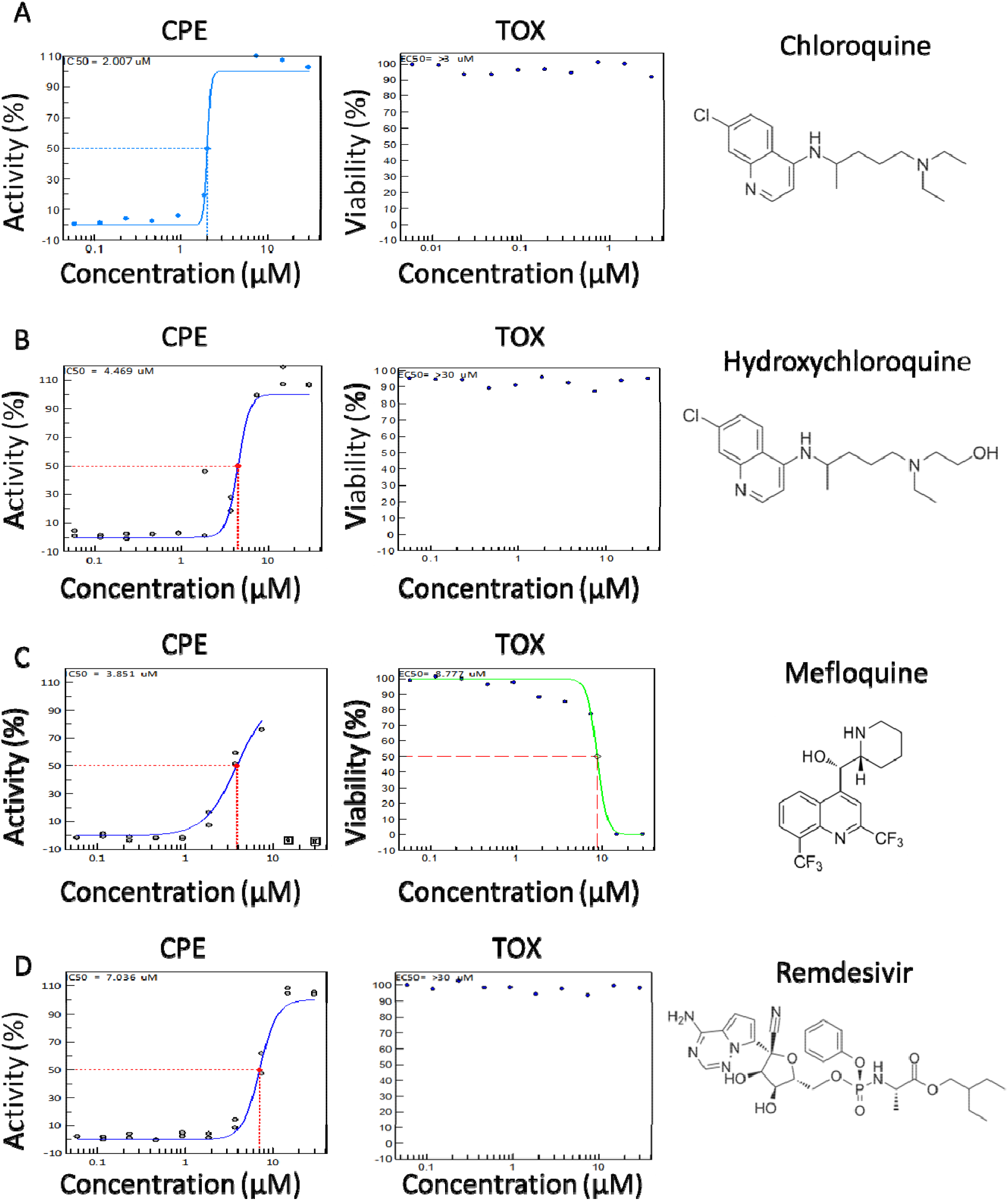
CPE activity and Toxicity for chloroquine, hydroxychloroquine, and mefloquine. **(A)** Chloroquine**, (B)** hydroxychloroquine, **(C)** mefloquine, and **(D)** remdesivir CPE activity (blue curve, left graph) and Toxicity (green curve, right graph) 10 point, 1:2 dilution concentration-response curves starting at 30.0 μM down to 2.29 nM, along with theirs structure. Dashed line indicates EC_50_ or CC_50_ for CPE and Toxicity assays, respectively. Duplicate values shown for each concentration. Curves generated using non-linear regression.

**Table 1.** CPE assay in Vero-E6 and average LC3B-based autophagy assay parameters from four cell lines

| Compound | MoA | CPE<br>EC<sub>50</sub><br>(μM) | CPE<br>CC<sub>50</sub><br>(μM) | CPE<br>SI | Autophagy<br>EC<sub>50</sub> (μM) | Autophagy<br>CC<sub>50</sub> (μM) | Autophagy<br>SI | MoA<br>Ref. |
| --- | --- | --- | --- | --- | --- | --- | --- | --- |
| Chloroquine | ↓ lysosome fusion | 2.01 | >30 | >10 | 3.29 ± 1.86 | >50 | >10 | (30) |
| ROC-325 | ↓ lysosome fusion | 3.28 | >30 | >10 | 5.2 ± 1.71 | >25 | >10 | (38) |
| Mefloquine | ↓ autophagic flux | 3.85 | 8.78 | 2.3 | 7.3* | 18.4 ± 2.08 | 2.6 | (46) |
| Hydroxy-chloroquine | ↓ lysosome fusion | 4.47 | >30 | >10 | 6.55 ± 6.67 | >50 | >10 | (12) |
| Hycanthone | ↑ lysosomal membrane permeabilization | 5.79 | 14.2 | 2.5 | 7.35 ± 4.7 | 11.3 ± 2.73† | 1.5 | (45) |
| Clomipramine | ↓ autophagic flux | 13.6 | >30 | >10 | 13.2 ± 5.4 | >50 | >10 | (42) |
| Verteporfin | ↓ autophagosome formation | ND | >30 | ND | ND | ND | ND | (75) |

\* Could only be calculated from Huh-7.5
† Max inhibition of cell viability ~60%
ND – not determined
SI>10 used when no CC<sub>50</sub> was calculated

### Autophagy inhibitors increase LC3B and LysoTracker dye staining

Because 6 out of the 7 autophagy inhibitors (ROC-325, clomipramine, hycanthone, CQ, HCQ, and mefloquine) showed activity in the CPE assay, we sought to confirm their effect on autophagy in Vero-E6, HeLa, HEK293T, and Huh-7.5 cells using immunostaining for autophagy marker LC3B, as well as LysoTracker dye staining. LC3B immunostaining directly visualizes autophagosomes, while LysoTracker Dye stains acidic organelles. These assays allow for the visualization of autophagosome accumulation and acidic organelles such as endosomes and lysosomes, respectively. Compounds that block autophagic flux are expected to increase LC3B and LysoTracker staining measurements (*34*).

To carry out this assay, cells were allowed to adhere overnight, and were then treated with compounds at concentrations ranging from 50 μM – 0.02 μM for approximately 16 h. In Vero-E6 cells, increases in intracellular LC3B spot, also called spots, were concentration-dependent for all of the compounds except for mefloquine (Fig. 4A,B). CQ, HCQ, and hycanthone treatment produced maximal spot counts, while ROC-325 and clomipramine produced a submaximal increase of 80% and 40%, respectively. Mefloquine was ineffective at inducing LC3B spot accumulation. Increases in LC3B spots indicate an accumulation of LC3B that is localized to the autophagosome when autophagic flux is blocked. The potent effect of CQ and HCQ on LC3B spot counts was apparent in all cell lines tested (Fig. 4 and Fig. S1,3,5). Based on nuclei counts, CQ, HCQ, clomipramine, and ROC-325 were not cytotoxic at the highest concentrations (50 μM for all except for ROC-325 at 25 μM). In line with the drug toxicity data from the CPE assay, mefloquine was completely toxic at 50 μM, while hycanthone killed approximately 60% of cells at 50 μM. The compound CC_50_ data was consistent between the two assays.

**Fig. 4.**
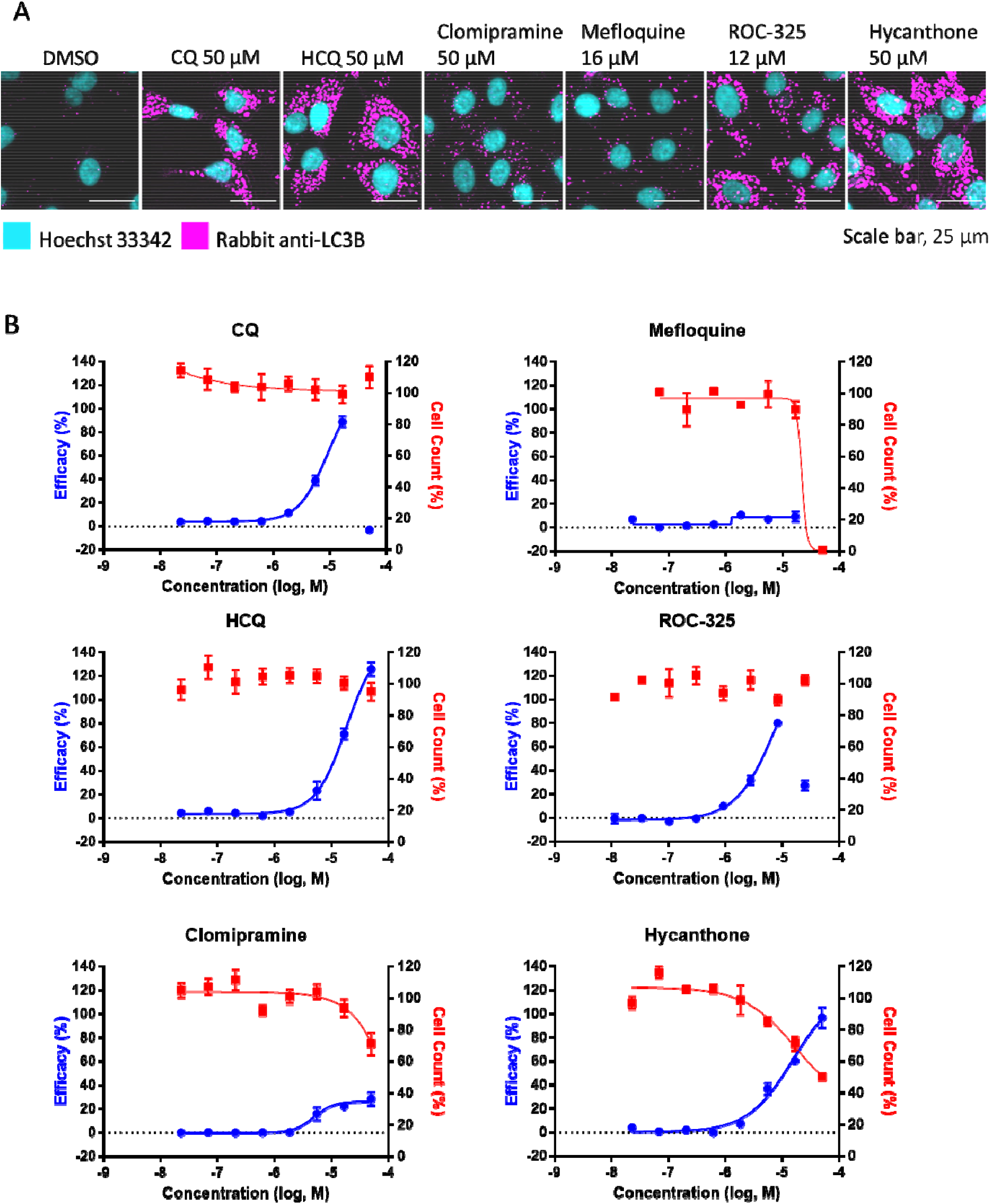
Autophagy inhibition assay using LC3B immunostaining in Vero-E6 cells. **(A)** Image montage of DMSO, CQ, HCQ, clomipramine, mefloquine, ROC-325, and hycanthone stained with Hoechst 33342 (cyan) and LC3B (magenta). CQ and HCQ images taken from wells in positive control column 2. Scale bar, 25 μm. **(B)** 8 point 1:3 dilution concentration-response curves starting at 50 μM down to 0.023 μM for compounds in A Blue curve indicates Efficacy, red curve indicates Cell Counts. Efficacy data normalized to DMSO (0%) and CQ (100%). Cell count data normalized to DMSO (100%) and 0 (no cells 0%). Error bars indicate SD. N = 3 intraplate replicates. Curves generated using non-linear regression.

In Vero-E6 cells, after drug treatment large concentration-dependent increases of LysoTracker relative spot intensity measurements were observed (Fig. 5A,B). With the exception of HCQ, the maximum efficacy was higher than the CQ positive control (100%) that was used to normalize the responses. Interestingly, clomipramine and mefloquine, which did not induce large increases in Vero-E6 LC3B spot counts, produced dramatic elevations in LysoTracker relative spot intensity similar to ROC-325 and hycanthone (Fig. 5B). In further support of the CPE assay data, mefloquine was toxic at the highest concentration.

**Fig. 5.**
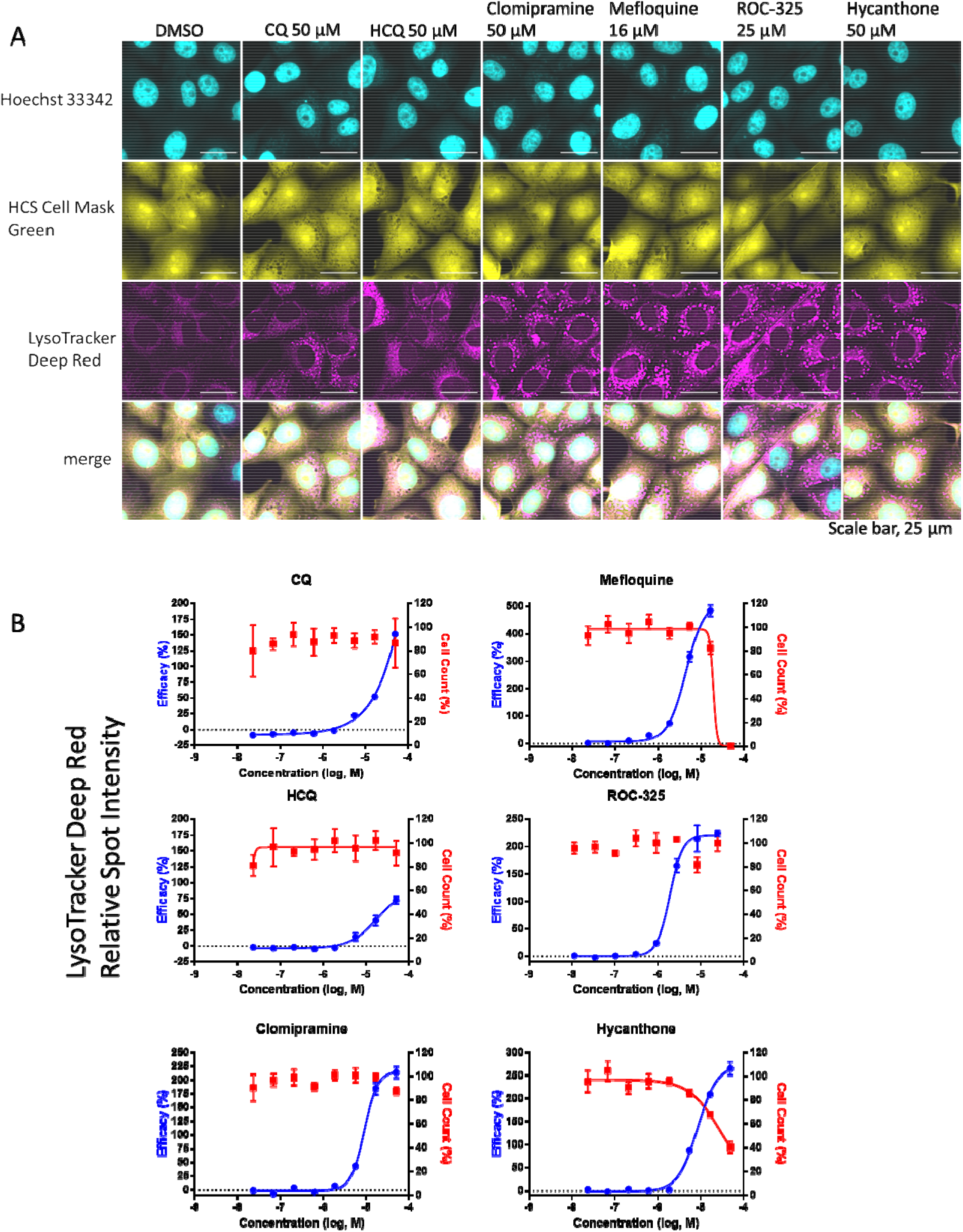
Autophagy inhibition assay using LysoTracker Deep Red staining in Vero-E6 cells. **(A)** Image montage of DMSO, CQ, HCQ, clomipramine, mefloquine, ROC-325, and hycanthone stained with Hoechst 33342 (cyan), HCS Cell Mask Green (yellow), and LysoTracker Deep Red (magenta). CQ and HCQ images taken from wells in positive control column 2. Scale bar, 25 μm. **(B)** 8 point 1:3 dilution concentration-response curves starting at 50 μM down to 0.023 μM for compounds in A. Blue curve indicates Efficacy, red curve indicates Cell Counts. Efficacy data normalized to DMSO (0%) and CQ (100%). Cell count data normalized to DMSO (100%) and 0 (no cells 0%). Error bars indicate SD. N = 3 intra-plate replicates. Curves generated using non-linear regression.

In addition to Vero-E6 cells, we also examined the effects of these compounds in three human cell lines and observed some differences between them (Fig. S1-6). For example, in Huh-7.5, mefloquine increased LC3B spot counts even at low concentrations (Fig. S3), whereas in other cell lines it was not a potent inducer of autophagosome accumulation. Clomipramine was effective in increasing LC3B in all cell lines except for Vero-E6 (Fig. 4, Fig. S1,3,5). In contrast, hycanthone and mefloquine produced the strongest effect on LysoTracker measurements in Vero-E6 compared to the other three cell lines (Fig. 5, Fig. S2,4,6). Although there were some interesting variations in compound effects among the cell lines tested, the average EC50 and CC50 values from the LC3B spot count measurements in all four cell lines corresponded well with the data from the CPE assay, indicating that the effects of the compounds on markers related to autophagy and protection from viral-induced cell death were well-correlated (Table 1).

We have illustrated our working hypothesis in Figure 6 as to one potential mechanism for the reduction of viral infection and subsequent CPE. First, in a healthy cell there is normal endocytosis of extracellular material and cellular components at the plasma membrane (Fig. 6A). When healthy cells are treated with autophagy inhibitors, the processes of endolysosome and autolysosome fusion are disrupted, leading to an increase in autophagosomes and late endosomes (Fig. 6B). In the case of an infected cell, potential endocytosis of SARS-CoV-2 leads to the release of viral RNA into the cell, whereas autophagic machinery may be hijacked to prevent flux (orange X) (Fig. 6C). We hypothesize that when autophagy inhibitors are present during viral infection, interference of multiple processes (red Xs) might lead to containment of the virus, and reduction in viral replication (Fig. 6D).

**Fig. 6.**
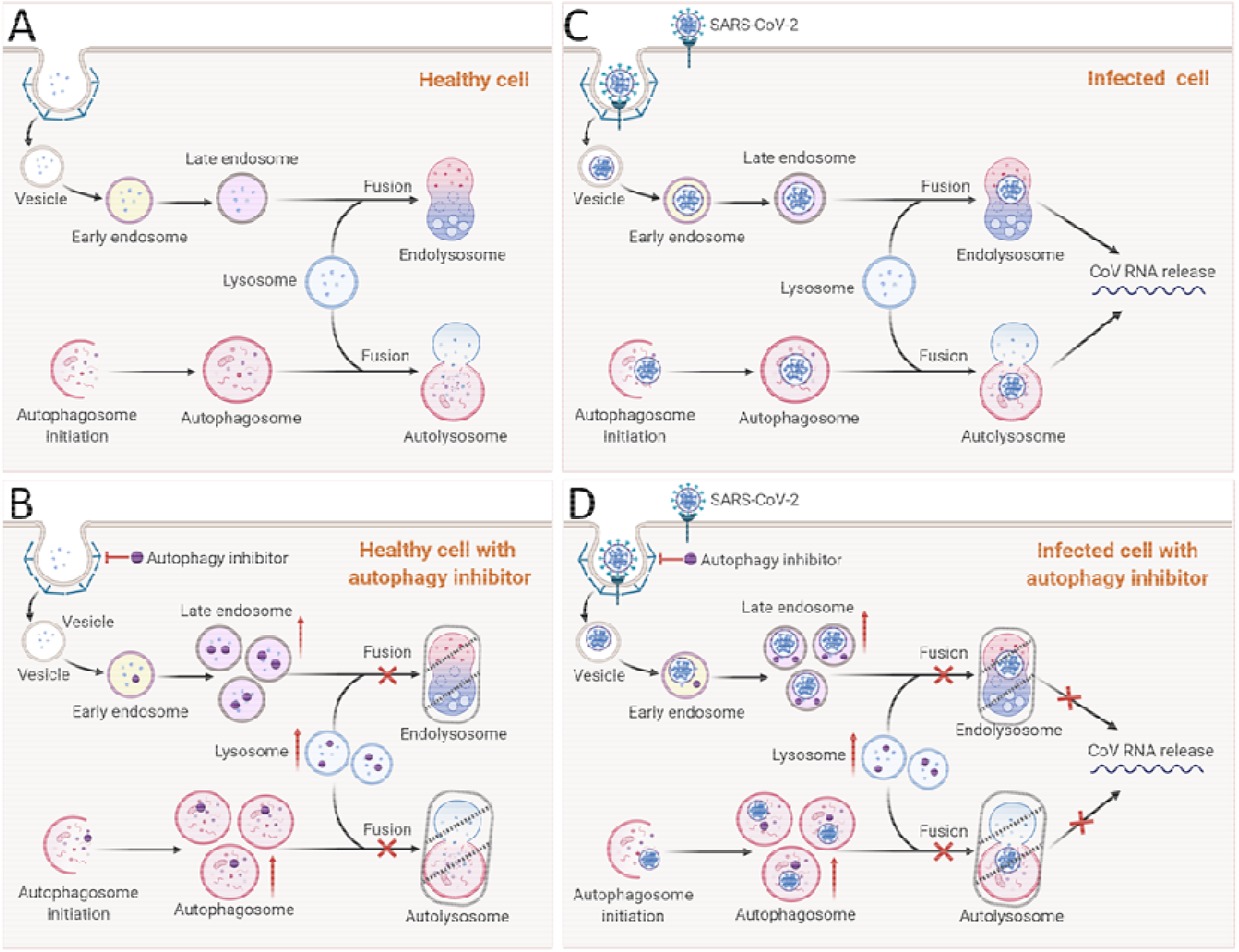
Illustration of autophagy inhibitors and their blockade of viral infection. (A) Healthy ce**lls** have normal autophagic flux and the endocytic pathway is functional. (B) Autophagy inhibitor treatment in healthy cells causes a blockade of normal fusion processes and a buildup of endosomes and autophagosomes. (C) In healthy cells, viral infection through endocytosis leads to the release of viral RNA after endosome lysosome fusion. Similarly, autophagy of viral particles may result in formation of viral autophagosomes but lysosome fusion would be blocked by the virus (orange X). Dotted arrow indicates a possible, but unverified event of viral RNA release from autophagosomes. (D) Autophagy inhibitors can block steps (red Xs) within the viral life cycle including at the early steps of endocytosis, the fusion of endosomes with the lysosome, to prevent the release of viral RNA and subsequent cell death.

## Discussion

New anti-viral drug repurposing opportunities are necessary for pre-clinical and clinical evaluation for treating COVID-19. In this work we have identified several autophagy inhibitors that can protect against CPE of SARS-CoV-2 in Vero-E6 cells. ROC-325 (*38–40*) and clomipramine (*41, 42*) display autophagy inhibitor activity that can completely prevent SARS-CoV-2 CPE without any significant inherent cytotoxicity. Hycanthone, an FDA-approved schistosomicide and oxidative metabolite of lucanthone (*43–45*), and mefloquine (*46–48*) both showed moderate levels of activity against SARS-CoV-2 CPE, but did exhibit drug-induced cell toxicity at the highest drug concentration tested (up to 30 μM). The autophagy inhibitor verteporfin, a benzoporphyrin derivative used in the clinic as a photosensitizer (*49*), did not inhibit CPE of SARS-CoV-2, and was not tested in follow-up autophagy assays. To confirm whether CPE protecting compounds interrupted cellular autophagy and lysosomal function, we examined their effects on autophagy marker LC3B (*50*), along with late endosome and lysosomes as visualized with LysoTracker dye. We found that the activities of autophagy inhibition as measured by LC3B spot counts correlated well with inhibition of SARS-CoV-2 measured in the CPE assay for ROC-325, clomipramine, hycanthone, and mefloquine. To our knowledge, this is the first report showing that ROC-325 and hycanthone are efficacious against SARS-CoV-2.

The 72 h SARS-CoV-2 CPE assay measures the phenotypic consequence of viral infection and replication in cells (*51–53*). SARS-CoV-2 can induce cell death (*54–57*) after 48 to 72 h of infection, and thus cell viability is a surrogate measure of viral replication *in vitro*. However, there are limitations to the CPE assay including its dependence on the host response and the fact that it is an indirect measurement of SARS-CoV-2 infection and replication. The phenotypic outcome can also vary depending on culture conditions and viral multiplicity of infection (MOI), number of virions that are added per cell during infection (*58*). The potencies of drug protection against virally-induced cell death can be lower than in other assays that directly measure viral load. Nevertheless, this study confirms that SARS-CoV-2 infection in Vero-E6 cells results in cell death similar to other reports, and that CPE can be suppressed by blocking autophagy with small molecule inhibitors to the same extent as positive control remdesivir (*59, 60*). Recently, a drug-repurposing screen of FDA-approved compounds, using a similar CPE assay with SARS-CoV-2 in Vero-E6 cells, found clomipramine (IC_50_ 5.93 μM; CC_50_ >30 μM) and mefloquine (IC_50_ 7.11 μM; CC_50_ >18.5 μM) to be active with low toxicity (*61*). The same study found HCQ to be more active than CQ with an IC_50_ of 9.21 μM and 42.03 μM, respectively. Mefloquine was also found to be active in another SARS-CoV-2 CPE screen using Caco-2 cells with an IC50 of 14.1 μM (*62*). In our study, the SI was calculated using the ratio of the EC50, the half-maximal effective concentration, and the CC50, the half-maximal cytotoxic concentration. Between the CPE and the autophagy assays there was good correspondence in the cytotoxicity measurements by CellTiter-Glo and nuclei counts, respectively. The SI is an important measure for future preclinical development, as it provides insights into the potential clinical safety of a compound at a cellular level. From this work, we show that CQ, HCQ, clomipramine, and ROC-325 were less than 50% cytotoxic at all concentrations, whereas mefloquine and hycanthone were cytotoxic at the highest concentrations with mefloquine being the most cytotoxic.

Evolution has endowed many viruses with the ability to escape autophagic degradation by using the autophagosome membrane for the formation of viral double membrane vesicles (DMVs), although the precise mechanism is still unclear. It has also been reported that some coronavirus proteins such as open reading frame protein 8b (ORF-8b), directly contribute to cell death following viral infection (*63*). Interestingly, ORF-8b causes the induction of autophagosome formation accompanied by damaging effects on lysosomal function and autophagy flux. ORF-8b also forms aggregates in cells that caused ER stress and lysosome malfunction, which could be responsible for reduced clearance of viral particles by autophagic flux (*63*). The nonstructural protein 6 (NSP-6) of the infectious bronchitis virus (IBV), an avian coronavirus, significantly increased the number of autophagosomes in host cells (*28*). The SARS-CoV accessory protein ORF-3a has three transmembrane domains that insert into the lysosomal membrane causing lysosome function dysregulation and necrotic cell death (*27*). Recently, Benvenuto and colleagues analyzed 351 available SARS-CoV-2 gene sequences and discussed that the mutations in NSP-6 may modify the virus’ activity for inducing autophagy, though experimental data was not presented (*21*). It appears paradoxical that viral infection inhibits autophagic clearance while autophagy inhibitors, also known to block autophagosome to lysosome fusion, suppress viral infection. Our data, combined with the reported mechanism of action for CQ as an antiviral, suggest that these autophagy inhibitors may interrupt the early steps in the viral life cycle, namely the fusion of the viral endosomes with the lysosome, thereby reducing viral replication and protecting cells from viral induced cell death. The effect of altering endosomal pH among other mechanisms appears to make compounds like HCQ and CQ highly effective against SARS-CoV-2 and other viruses (*64*). However, more work is needed to elucidate the exact mechanism of action for these autophagy inhibitors in relation to SARS-CoV-2 and the impact at the different stages of the viral life cycle. Other host targets for viral inhibition include the point of entry with clathrin-mediated endocytosis of the virus (*65*), p38 MAPK involved in viral replication (*66*), post-translational processing of viral proteins in the Golgi apparatus (*67*), and budding of the virus from the infected cell (*68, 69*).

ROC-325 was originally developed as an orally available inhibitor of autophagy designed to incorporate the chemical motifs of HCQ and lucanthone, with the goal of both improved autophagic inhibition and consequent single-agent anticancer activity (*39, 45*). ROC-325 is a preclinical candidate with low *in vitro* and *in vivo* toxicity and strong anti-cancer properties (*40, 70*). Our study shows that it may also be a candidate for repositioning as a treatment for COVID-19. Clomipramine, a centrally acting, FDA-approved, tricyclic antidepressant used for the treatment of obsessive-compulsive disorder, panic disorder, major depressive disorder, and chronic pain (*71, 72*) may also be an interesting preclinical candidate with its existing regulatory status easing a path towards use in the clinic, although the human Cmax does not cover the CPE EC50. Because most of these compound EC50 values were higher than their human plasma concentrations at the clinically efficacious doses, they likely will not be efficacious as single agents for the treatment of COVID-19 (Table 2). Indeed, toxicity with CQ and HCQ has been reported and caution has to be taken with its clinical application because of potential cardiotoxicity (*73*). Furthermore, a large observational trial did not find a reduction in death of patients taking HCQ, which suggests that large randomized clinical trials are needed to assess the true benefit to patients with regard to decreased mortality rate and duration of hospitalization (*74*). However, the sum of this work indicates that targeting steps of the viral life cycle in cells with molecules similar to CQ, focusing on their anti-autophagic properties, could be a valid drug discovery strategy for combating SARS-CoV-2. The compounds described here also have value as research tools to better understand the interplay between host autophagy pathway and viral live cycle.

**Table 2.**
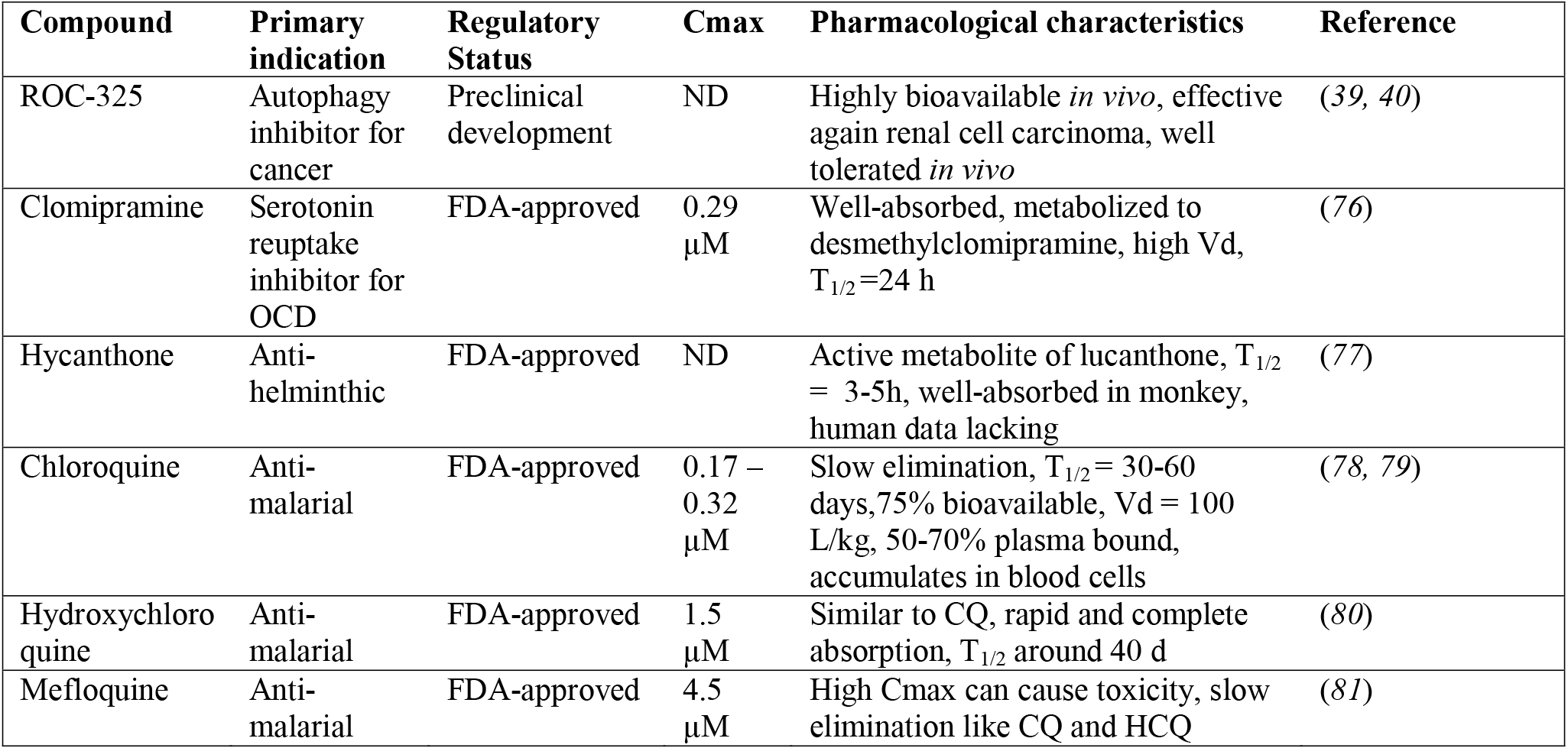
Clinical features of autophagy inhibitor compounds

Because such compounds target host cells to suppress SARS-CoV-2 CPE, they have potential to be combined with other drugs that directly target viral proteins for treatment. This type of combination therapy has certain advantages including synergistic activity from different mechanisms of action and reducing the development of viral drug resistance due to the involvement of a host cell target. Furthermore, individual drug concentrations can be lowered in combination therapies to prevent the toxicity seen at higher doses when treating with a single drug. Further tests of the drug combination therapy using SARS-CoV-2 animal models will be needed to confirm the therapeutic usage of these compounds.

## Data and materials availability

Data is available upon request.

## Materials and Methods

### Study Design

#### Sample Size and Replicates

For the CPE, inter-plate duplicates were used for each data point for quantitative HTS and curve fitting. For CPE luminescence measurements, each well was read once. For the autophagy assay, three intra-plate replicates were used in consecutive columns for quantitative HTS, high-content analysis, and curve fitting. For the autophagy assay automated high-content imaging, each well was imaged 6 times in equally spaced fields using a 40x objective. This allowed for the collection of data from approximately 500 or more cells per well.

#### Data inclusion and outliers

All data was included in the CPE assay. For the high-content imaging autophagy assays, efficacy data points were excluded in the case where there was >80% cell death. For non-linear curve fitting, data points were excluded when there was an experimental error that prevented proper drug addition or staining.

#### Selection of endpoints

CPE assay 72 h and autophagy assay 16-18 h endpoints were selected *a priori* based on previous studies.

#### Research Objectives

We aimed to contribute valuable pharmacological data towards the fight against COVID-19 by screening autophagy inhibitor compounds in a viral cytopathic effect assay to determine their potency and efficacy in preventing virally-induced cell death. We further aimed to validate these autophagy inhibitors in a number of cell lines to understand whether the pharmacological effect of autophagy inhibition corresponded with anti-viral effects. Autophagy inhibition is a known anti-viral strategy effective *in vitro*, *in vivo*, and potentially in human patients. However, there is a lack of clinically available autophagy inhibitors due to dose-limiting adverse side effects. After screening, we identified a new preclinical compound ROC-325 is a potential target for further development.

#### Units of investigation

Traditional cell culture methods were used in high-throughput formats for CPE and autophagy screening. Vero-E6 cells previously developed by collaborators were used for the CPE assay. Vero-E6, HEK293T, and HeLa cells were purchased from ATCC, and Huh-7.5 cells were a gift from the Tang Lab at FSU.

#### Experimental design

The experiments put forth in this research article were controlled laboratory experiments devised with the guidelines established for high-throughput screening. Cells were maintained in a healthy state with proper cell culture techniques and treated using small volumes of compound dissolved in DMSO. Luminescence readings were collected for the CPE assay and fluorescence images were captured using automated high-content microscopy. Controls were assigned to specific wells and compounds were distributed throughout the entire 384 well plates. For the autophagy assays, compound dilutions were arranged vertically with the highest concentration in the middle of the plate and the lowest concentrations on the edges. Each compound was in three consecutive columns. Further details are provided in the Methods section. Measurements for the CPE assay and fluorescence images were captured sequentially well by well. For the autophagy assay, a horizontal serpentine imaging sequence was used.

#### Blinding

For the luminescence readings, the simple data structure was processed according to the plate layout annotation. For the autophagy assay, a custom high-content imaging protocol was developed in Columbus Analyzer for each cell line based on the detection of signals from the controls and the processing was automated. The data was initially processed using compound identifiers called NCGC values, and then the data was quantified and visualized in Excel and Prism GraphPad. The compound NCGC numbers were then unmasked using the corresponding compound names.

### Reagents

The following items were purchased from Gibco: MEM (11095), DMEM (11965092), HI FBS (14000), Pen/Strep (15140). TrypLE (12604013), PBS -/- (w/o Ca^2+^ or Mg^2+^) (10010049), Trypsin-EDTA (25300-054). Hyclone FBS (SH30071.03) was purchased from GE Healthcare. The following items were purchased from ATCC: EMEM (30-2003), Vero-E6 (CRL-1586, RRID:CVCL_0574), HeLa (CCL-2, RRID:CVCL_0030), HEK293T (CRL-3216, RRID:CVCL_0063). Huh-7.5 cells were a gift from the Tang Lab at FSU. The following items were purchased from Invitrogen: Live Cell Imaging Buffer (A14291DJ), LysoTracker Deep Red (L12492), goat-anti-mouse AlexaFluor-647 (A-21242, RRID:AB_2535811), HCS Cell Mask Green (H32714), Hoechst 33342 (H3570). LC3B primary rabbit antibody (3868S, RRID:AB_2137707) was purchased from Cell Signaling Technologies. Cell Staining Buffer (420201) was purchased from BioLegend. The following items were purchased from Corning: 384-well plates (3764 BC), BioCoat 384-well poly-d-Lysine coated plates (354663 BC), Amphotericin B (30-003-CF). 100% Methanol (34860) was purchased from Sigma-Aldrich. Calpain Inhibitor IV (208724) was purchased from CalbioChem.

### Cell Culture

Vero-E6 cells previously selected for high ACE2 expression (*82*) were cultured in MEM/10% HI FBS supplemented with 0.5 μg/mL amphotericin B and passaged twice per week at 1:5 dilutions using trypsin. Briefly, cell culture media was aspirated, and cells were washed twice with PBS. 2 mL of trypsin is added for 1-2 minutes at room temperature and 10 mL of EMEM is added to wash flask and create a single cell suspension. Cells are spun at 800 RPM for 5 minutes. Supernatant was aspirated and cells resuspended in fresh media for seeding into flasks or multi-well plates.

Vero-E6 (grown in EMEM, 10% FBS, and 1% Penicillin/Streptomycin), HeLa CCL-2, HEK293T and Huh-7.5 (grown in DMEM, 10% FBS, and 1% Penicillin/Streptomycin) were cultured in T175 flasks and passaged at 95% confluency. Briefly, cells were washed once with PBS and dissociated from the flask using TrypLE. Cells were counted prior to seeding.

### Preparation of Assay Ready Plates

An 80 μL aliquot of each compound stock solution (10 mM in 100% DMSO) is transferred into empty wells in columns 3 and 13 of an Echo^®^ Qualified 384-Well Polypropylene Source Microplate (Labcyte P-05525). Compounds are diluted 2-fold by transferring 40 μL of each sample into 40 μL DMSO in the adjacent well (columns 4 and 14) and mixing. This process is repeated to create 8 more 2-fold serially diluted samples in the wells of columns 5-12 and 6-22. Using a Labcyte ECHO 550 (San Jose, CA) acoustic liquid handling system a 90 nL aliquot of each diluted sample is dispensed into corresponding wells of a Corning 3764BC plate. An equal volume of DMSO is added to control wells to maintain 0.3% DMSO final assay concentration in all wells. These are referred to as Assay Ready Plates (ARPs) and are stored at −20°C.

### Method for measuring antiviral effect of compounds

A CPE assay previously used to measure antiviral effects against SARS-CoV (*35*) was adapted for performance in 384 well plates to measure CPE of SARS CoV-2 with the following modifications. Cells harvested and suspended at 160,000 cells/ml in MEM/1% PSG/1% HEPES supplemented 2% HI FBS were batch inoculated with SARS CoV-2 (USA_WA1/2020) at M.O.I. of approximately 0.002 which resulted in approximately 5% cell viability 72 h post infection. ARPs were brought to room temperature and 5μl of assay media was dispensed to all wells. The plates were transported into the BSL-3 facility were a 25 μL aliquot of virus inoculated cells (4000 Vero E6 cells/well) was added to each well in columns 3-24. The wells in columns 23-24 contained virus infected cells only (no compound treatment). A 25 μL aliquot of uninfected cells was added to columns 1-2 of each plate for the cell only (no virus) controls. After incubating plates at 37°C with 5% CO_2_ and 90% humidity for 72 h, 30 μL of Cell Titer-Glo (Promega, Madison, WI) was added to each well. Following incubation at room temperature for 10 minutes the plates were sealed with a clear cover and surface decontaminated and luminescence was read using a Perkin Elmer Envision (Waltham, MA) plate reader to measure cell viability. Raw data from each test well was normalized to the average signal of non-infected cells (Avg. Cells; 100% inhibition) and virus infected cells only (Avg. Virus; 0% inhibition) to calculate % inhibition of CPE using the following formula: % inhibition CPE = 100*(Test Cmpd – Avg. Virus)/(Avg. Cells – Avg. Virus).

### Method for measuring cytotoxic effect of compounds

Compound cytotoxicity was assessed in a BSL-2 counter screen as follows: Host cells in media were added in 25 μl aliquots (4000 cells/well) to each well of assay ready plates prepared with test compounds as above. Cells only (100% viability) and cells treated with hyamine at 100 μM final concentration (0% viability) serve as the high and low signal controls, respectively, for cytotoxic effect in the assay. DMSO was maintained at a constant concentration for all wells (0.3%) as dictated by the dilution factor of stock test compound concentrations. After incubating plates at 37°C/5% CO_2_ and 90% humidity for 72 h, plates were brought to room temperature and 30μl Cell Titer-Glo (Promega) was added to each well. Luminescence was read using a BMG PHERAstar plate reader following incubation at room temperature for 10 minutes to measure cell viability.

### Autophagy assays

20 μL of cells were seeded into 384-well, black, clear-bottom, poly-d-lysine coated plates to achieve 60% confluent wells. Plates were covered with metal lids and placed in a 37°C incubator with 95% humidity and 5% CO2 overnight before compound treatment. 100 nL of compound per well was dispensed using the Labcyte Echo 655. The compounds were added at 8 concentrations with 1:3 dilutions starting at 50 μM down to 0.02 μM. ROC-325 was dispensed at the highest working concentration of 25 μM due to a maximum solubility of 5 mM in DMSO.

For LysoTracker staining, 5 μL of a 5x 250 nM LysoTracker Deep Red (Invitrogen, Carlsbad, CA) diluted in Live Cell Imaging Buffer (Invitrogen) was added to the plates mentioned above and incubated for 1 h at 37°C with 5% CO2 after which cells were fixed using 4% PFA (Electron Microscopy Sciences, Hatfield, PA) and incubated at room temperature for 30 minutes. Media in wells was then evacuated and cells were washed three times with PBS using the automated Bluewasher plate washing system from Blue Cat Bio (Concord, MA). Plates were then sealed and imaged on the IN Cell 2500 HS (GE Healthcare, Chicago, IL) automated high-content imaging system. Images was uploaded to Columbus Analyzer and processed for high-content analysis.

For LC3B immunostaining, media was evacuated on the Bluewasher and 100% ice-cold methanol was added to wells for 10 minutes at −30°C. Plates were washed three times with PBS and blocked with Cell Staining Buffer (BioLegend, San Diego, CA). Plates were then incubated with rabbit-anti-LC3B (Cell Signaling Technologies, Danvers, MA) antibodies in Cell Staining Buffer for 2 h at room temperature. Plates were washed three times with PBS and secondary antibody goat-anti-mouse AlexaFluor-647 (Invitrogen) were added in Cell Staining Buffer for 1 h. Plates were washed three times in PBS before adding 1:5000 Hoechst 33342 (Invitrogen). After a final three washes in PBS, plates were sealed and imaged on the IN Cell 2500 HS automated high-content imaging system. Images was uploaded to Columbus Analyzer and processed for high-content analysis. Image montages were prepared using Fiji (ImageJ, NIH).

### Statistical analysis

CPE assay raw data from each test well was normalized to the average signal of non-infected cells (Avg. Cells; 100% inhibition) and virus infected cells only (Avg. Virus; 0% inhibition) to calculate % inhibition of CPE using the following formula: % inhibition CPE = 100*(Test Cmpd – Avg Virus)/(Avg Cells - Avg Virus). EC50 values were obtained using non-linear regression.

High-content image analysis data was downloaded as a Microsoft Excel spreadsheet. DMSO negative control (0% activity) (col. 1 and 24 for acoustic dispensing) and CQ positive control (100% activity) (col. 2 for acoustic dispensing, 8 wells) was used to normalize each compound concentrations’ response. The other 8 wells of column 2 contained 10 mM HCQ. EC50 values were obtained using non-linear regression in Graphpad Prism 7.04. In some cases, the highest concentration point was not included in curve-fit due to technical issues during experimental execution, although the measured value was shown. When cell viability was below 20%, the efficacy point was excluded altogether (i.e. mefloquine at 24 μM or 8 μM). Six fields per well were imaged on the IN Cell 2500HS. LC3B and LysoTracker data was obtained using a single well with hundreds of cells for each compound concentration from three intra-plate replicate wells were imaged when acoustic dispensing was used for compound treatment. Cell counts were also reported using nuclear object segmentation. GraphPad Prism 7.04v was used for visualizing autophagy data. EC50 and CC50 values from high-content imaging were obtained using non-linear regression.

## Ancillary information

### • Supporting Information

- Fig. S1. Autophagy inhibition assay using LC3B immunostaining in HeLa cells
- Fig. S2. Autophagy inhibition assay using LC3B immunostaining in HEK293T cells
- Fig. S3. Autophagy inhibition assay using LC3B immunostaining in Huh-7.5 cells
- Fig. S4. Autophagy inhibition assay using LysoTracker Deep Red staining in HeLa cells
- Fig. S5. Autophagy inhibition assay using LysoTracker Deep Red staining in HEK293T cells
- Fig. S6. Autophagy inhibition assay using LysoTracker Deep Red staining in Huh-7.5 cells

### • Corresponding Author Information

- Kirill Gorshkov –
- Wei Zheng –

### • Author Contributions

- Experimental – KG, CC, RB, LR, MX, MP
- Data Analysis – KG, BN
- Compound Management & Selection – KG, XH, PS, ZI
- Manuscript Conception – KG, CC, WZheng
- Manuscript Writing – KG, CC, KS, WH, WZheng
- Preparation of Figures – KG, CC, WZhu
- Critical Manuscript Editing and Discussion– KG, MP, CC, WZheng, AS, MH, MS, DL
- Laboratory Automation – CK-T, SM

## • Acknowledgment

- We thank Richard Eastman (NCATS) and Sara McKellip (SRI) for assistance with acoustic dispensing support and compound management. We thank the laboratory of Dr. Hengli Tang for providing the Huh-7.5 cells.

## • Abbreviations Used

SARS-CoV-2: Severe acute respiratory syndrome coronavirus 2
COVID-19: coronavirus disease that was discovered in 2019
CQ: chlorquine
HCQ: hydroxychloroquine
LC3B: microtubule-associated proteins 1B light chains 3B
CPE: cytopathic effect
SI: selectivity index
TOX: toxicity measurement for CC50 calculation
MOI: multiplicity of infection

## Supplementary Materials

**Fig. S1.**
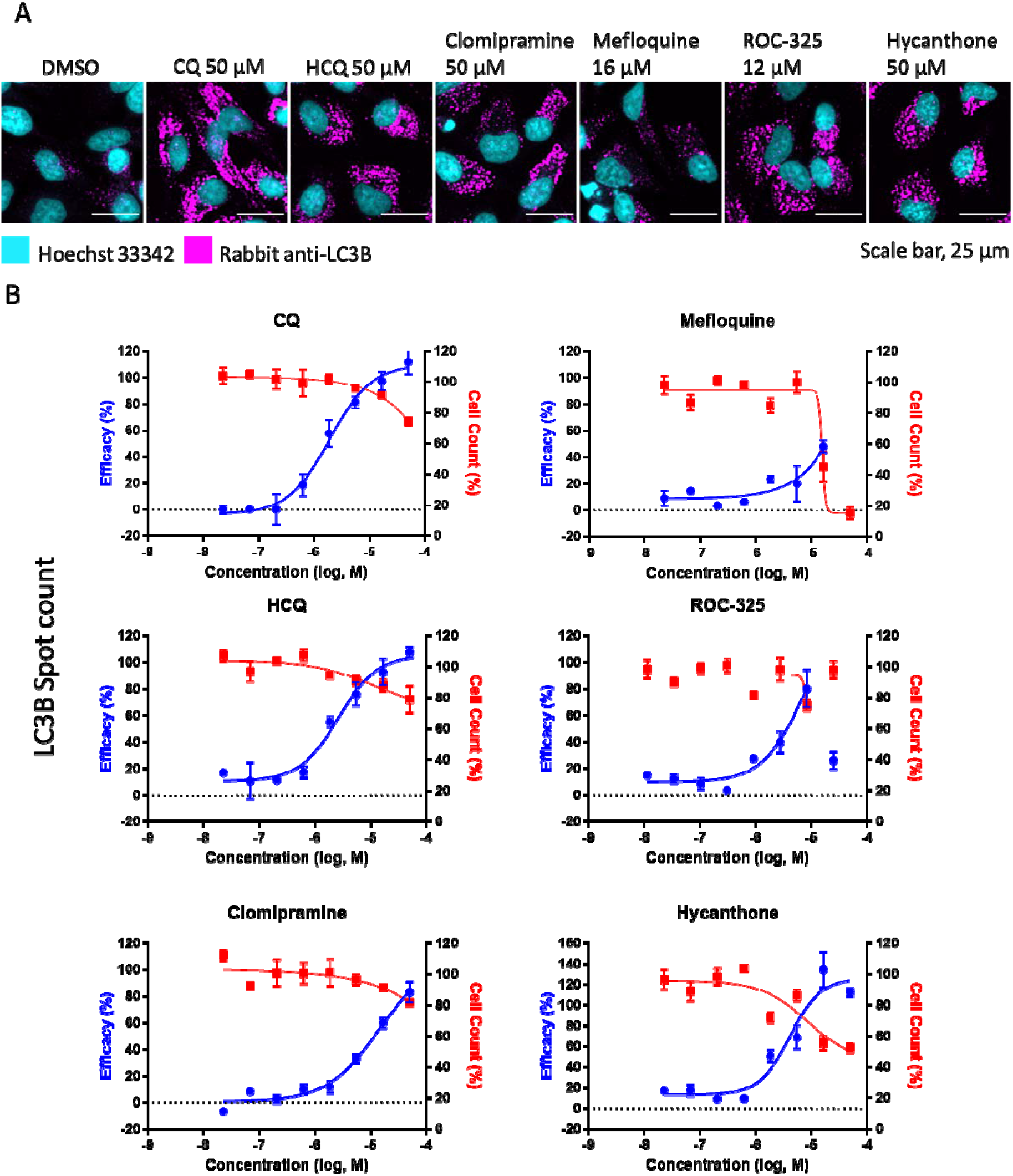
Autophagy inhibition assay using LC3B immunostaining in HeLa cells. **(A)** Image montage of DMSO, CQ, HCQ, clomipramine, mefloquine, ROC-325, and hycanthone stained with Hoechst 33342 (cyan) and LC3B (magenta). CQ and HCQ images taken from wells in positive control column 2. Scale bar, 25 μm. **(B)** 8 point 1:3 dilution concentration-response curves starting at 50 μM down to 0.023 μM for compounds in A. Blue curve indicates Efficacy, red curve indicates Cell Counts. Efficacy data normalized to DMSO (0%) and CQ (100%). Cell count data normalized to DMSO (100%) and 0 (no cells 0%). Error bars indicate SD. N = 3 intraplate replicates. Curves generated using non-linear regression.

**Fig. S2.**
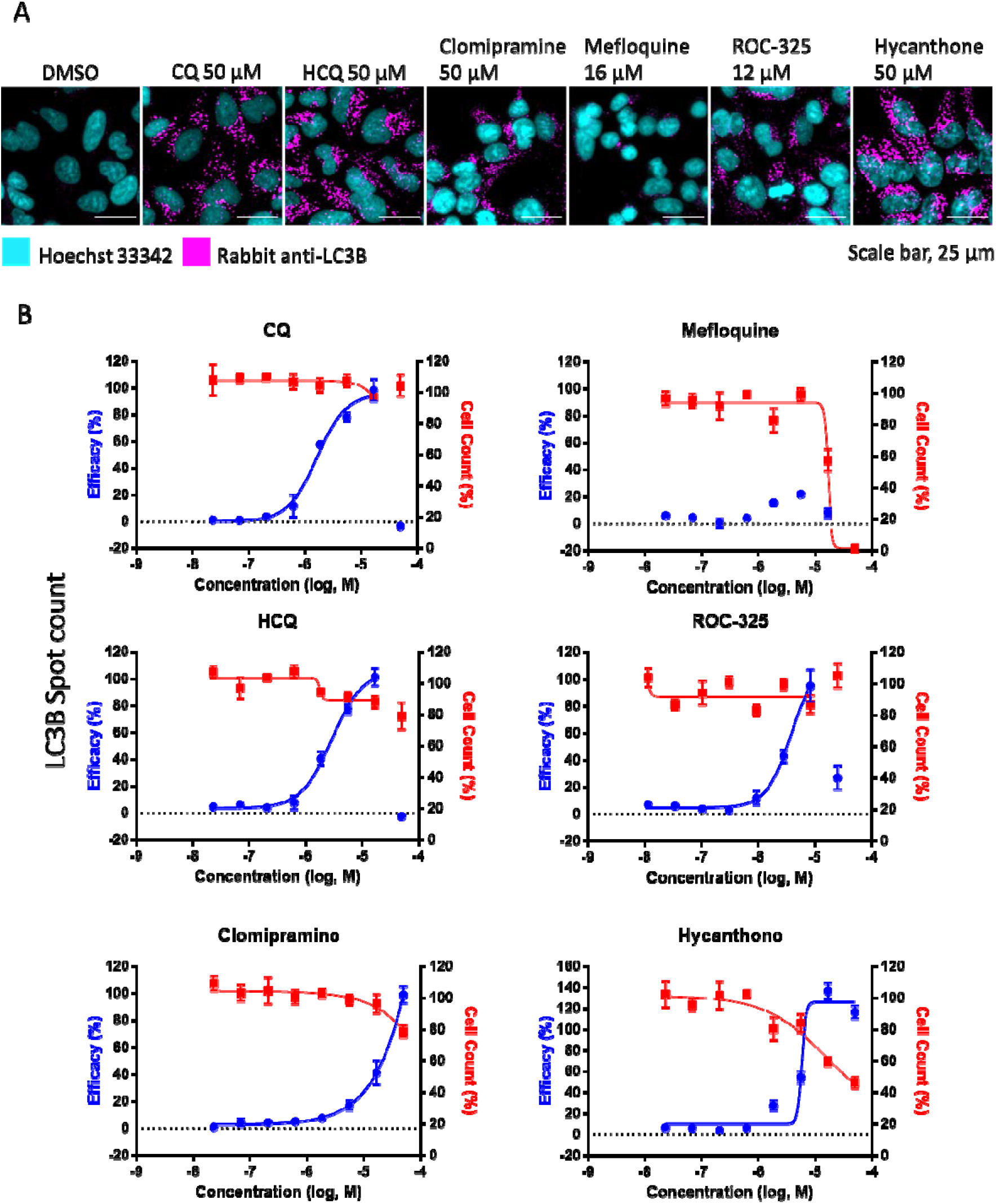
Autophagy inhibition assay using LC3B immunostaining in HEK293T cells. **A)** Image montage of DMSO, CQ, HCQ, clomipramine, mefloquine, ROC-325, and hycanthone stained with Hoechst 33342 (cyan) and LC3B (magenta). CQ and HCQ images taken from wells in positive control column 2. Scale bar, 25 μm. **(B)** 8 point 1:3 dilution concentration-response curves starting at 50 μM down to 0.023 μM for compounds in A. Blue curve indicates Efficacy, red curve indicates Cell Counts. Efficacy data normalized to DMSO (0%) and CQ (100%). Cell count data normalized to DMSO (100%) and 0 (no cells 0%). Error bars indicate SD. N = 3 intraplate replicates. Curves generated using non-linear regression.

**Fig. S3.**
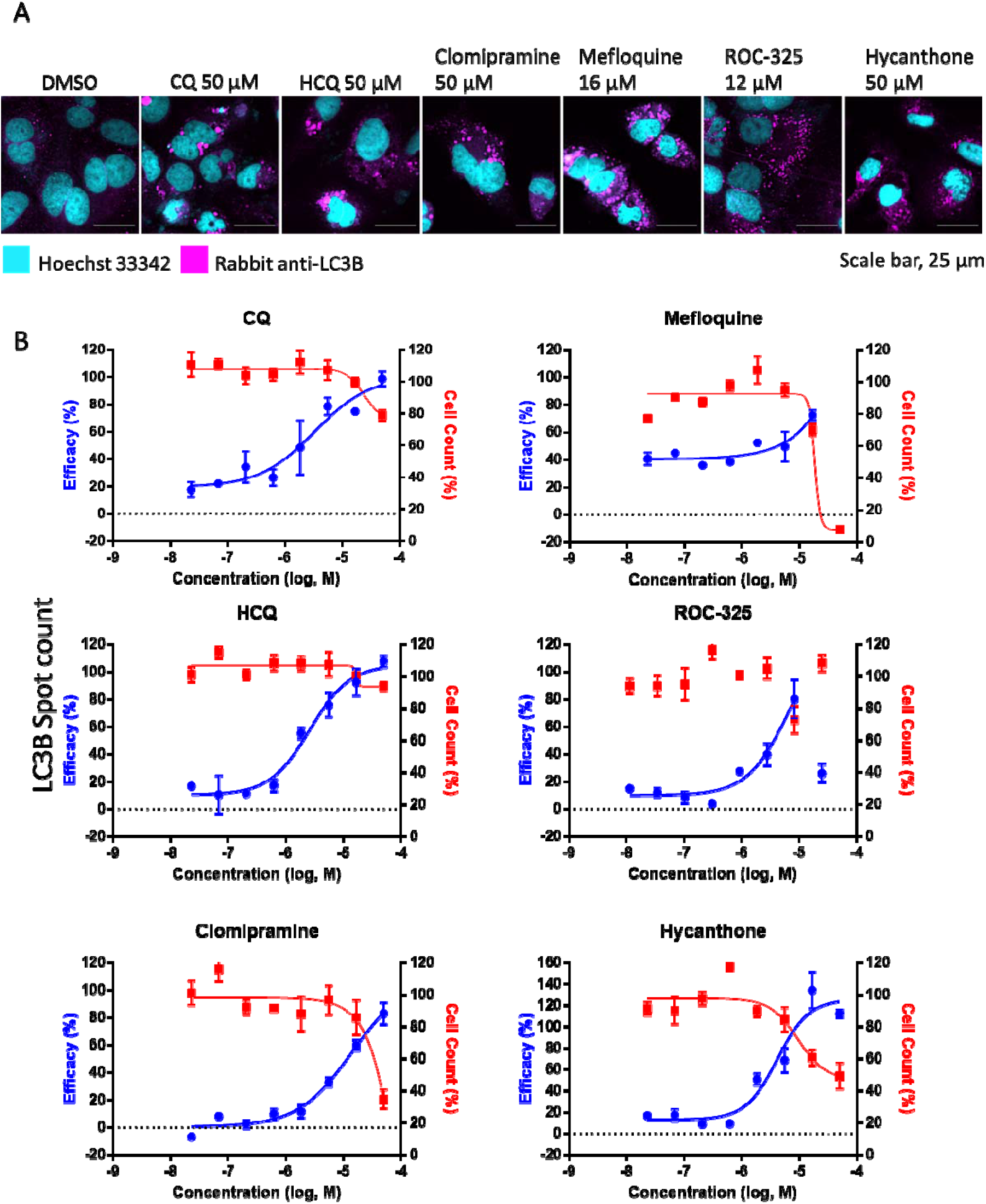
Autophagy inhibition assay using LC3B immunostaining in Huh-7.5 cells. **(A)** Image montage of DMSO, CQ, HCQ, clomipramine, mefloquine, ROC-325, and hycanthone stained with Hoechst 33342 (cyan) and LC3B (magenta). CQ and HCQ images taken from wells in positive control column 2. Scale bar, 25 μm. **(B)** 8 point 1:3 dilution concentration-response curves starting at 50 μM down to 0.023 μM for compounds in A. Blue curve indicates Efficacy, red curve indicates Cell Counts. Efficacy data normalized to DMSO (0%) and CQ (100%). Cell count data normalized to DMSO (100%) and 0 (no cells 0%). Error bars indicate SD. N = 3 intraplate replicates. Curves generated using non-linear regression.

**Fig. S4.**
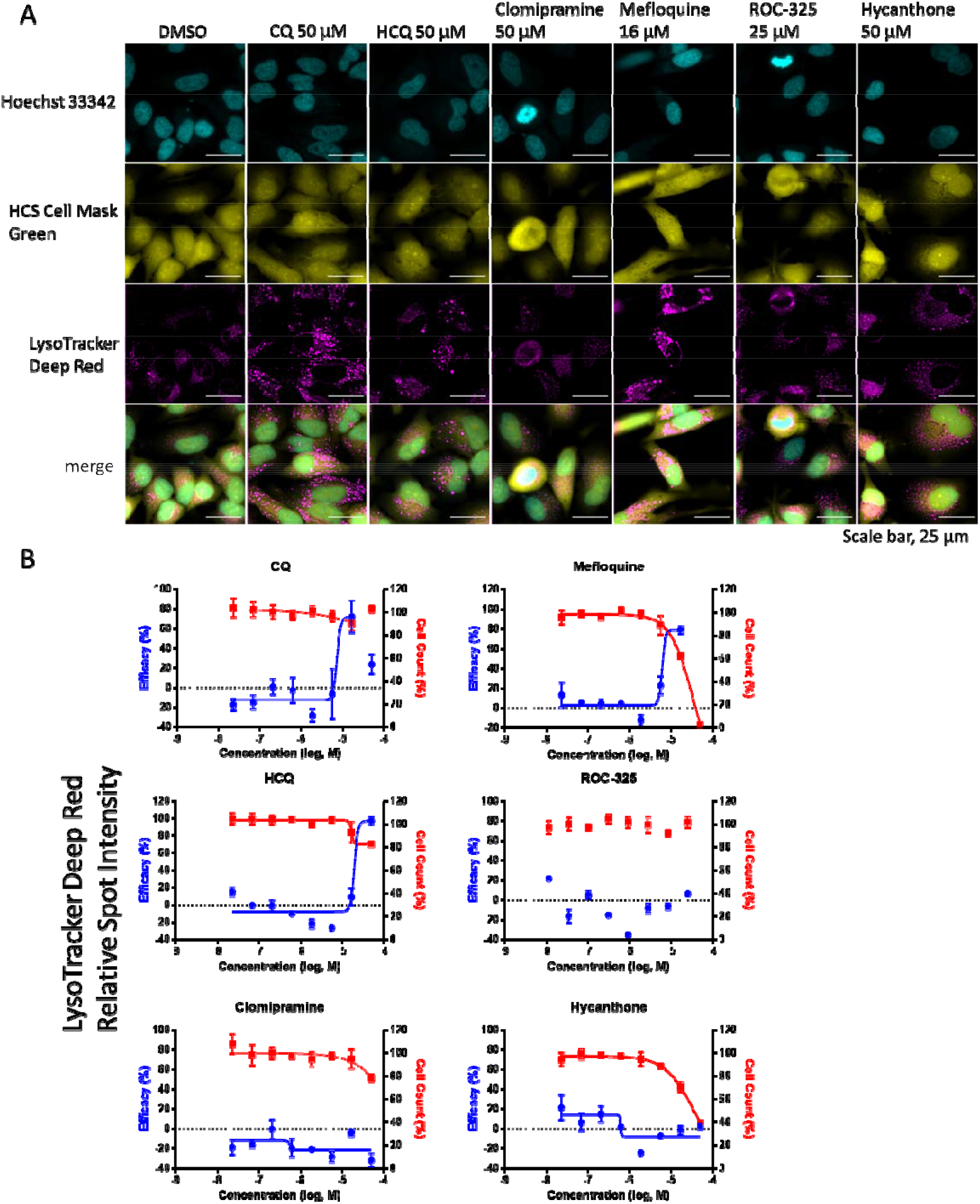
Autophagy inhibition assay using LysoTracker Deep Red staining in HeLa cells. **(A)** Image montage of DMSO, CQ, HCQ, clomipramine, mefloquine, ROC-325, and hycanthone stained with Hoechst 33342 (cyan), HCS Cell Mask Green (yellow), and LysoTracker Deep Red (magenta). CQ and HCQ images taken from wells in positive control column 2. Scale bar, 25 μm. **(B)** 8 point 1:3 dilution concentration-response curves starting at 50 μM down to 0.023 μM for compounds in A. Blue curve indicates Efficacy, red curve indicates Cell Counts. Efficacy data normalized to DMSO (0%) and CQ (100%). Cell count data normalized to DMSO (100%) and 0 (no cells 0%). Error bars indicate SD. N = 3 intra-plate replicates. Curves generated using non-linear regression.

**Fig S5.**
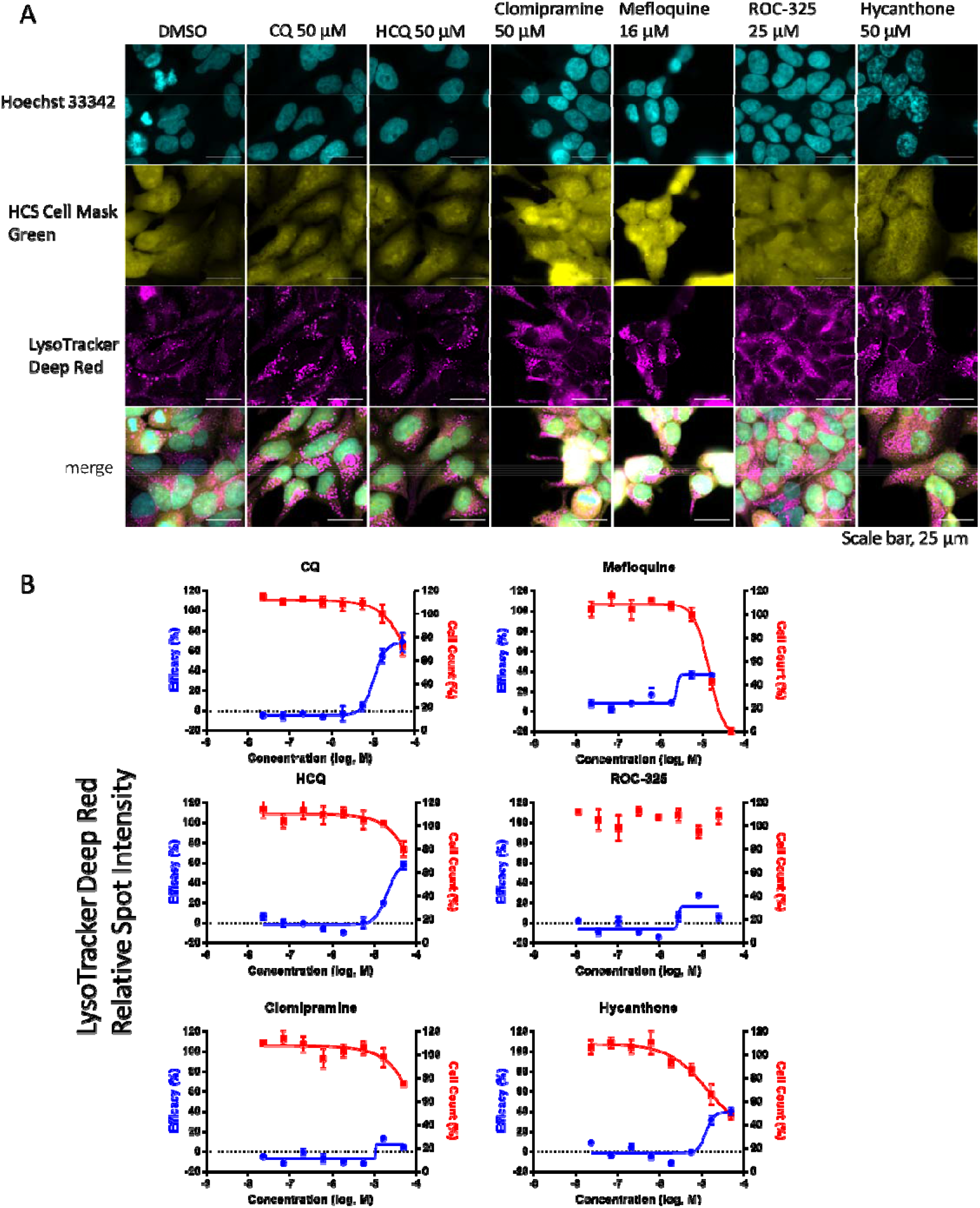
Autophagy inhibition assay using LysoTracker Deep Red staining in HEK293T cells. **(A)** Image montage of DMSO, CQ, HCQ, clomipramine, mefloquine, ROC-325, and hycanthone stained with Hoechst 33342 (cyan), HCS Cell Mask Green (yellow), and LysoTracker Deep Red (magenta). CQ and HCQ images taken from wells in positive control column 2. Scale bar, 25 μm. **(B)** 8 point 1:3 dilution concentration-response curves starting at 50 μM down to 0.023 μM for compounds in A. Blue curve indicates Efficacy, red curve indicates Cell Counts. Efficacy data normalized to DMSO (0%) and CQ (100%). Cell count data normalized to DMSO (100%) and 0 (no cells 0%). Error bars indicate SD. N = 3 intra-plate replicates. Curves generated using non-linear regression.

**Fig S6.**
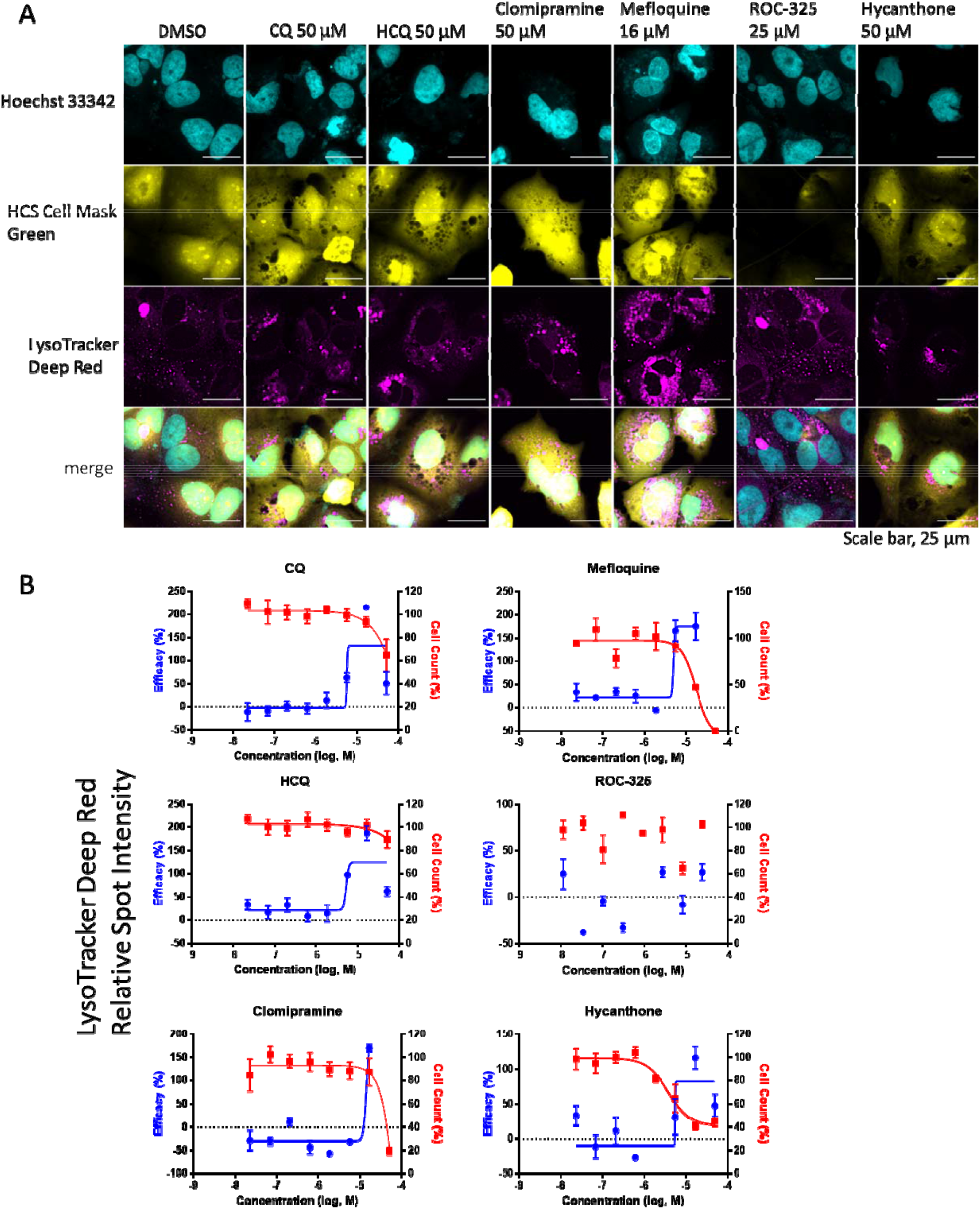
Autophagy inhibition assay using LysoTracker Deep Red staining in Huh-7.5 cells. **(A)** Image montage of DMSO, CQ, HCQ, clomipramine, mefloquine, ROC-325, and hycanthone stained with Hoechst 33342 (cyan), HCS Cell Mask Green (yellow), and LysoTracker Deep Red (magenta). CQ and HCQ images taken from wells in positive control column 2. Scale bar, 25 μm. **(B)** 8 point 1:3 dilution concentration-response curves starting at 50 μM down to 0.023 μM nM for compounds in A. Blue curve indicates Efficacy, red curve indicates Cell Counts. Efficacy data normalized to DMSO (0%) and CQ (100%). Cell count data normalized to DMSO (100%) and 0 (no cells 0%). Error bars indicate SD. N = 3 intra-plate replicates. Curves generated using non-linear regression.

